# Executive Functions in relation to Autonomic Control: An Overview of Neuropsychological Evaluation Methods

**DOI:** 10.1101/2025.02.25.640212

**Authors:** Shyama Shah, Janki Kuber, Gregory F. Lewis

## Abstract

Executive functions are a set of cognitive processes essential for cognitive control and coordination, enabling the achievement of objectives. These functions include mental exploration of ideas, reasoning, logical conclusion drawing, discipline, decision making, thoughtful consideration before action, tackling unforeseen challenges, resisting temptations, and maintaining focus. Numerous neuropsychological tests assess executive functions in relation to different autonomic control pathways that regulate involuntary physiological processes.

The complexity of the autonomic nervous system and the challenges in measuring it alongside cognitive assessments are significant. Issues such as subject movement, environmental changes, and time-consuming protocols further complicate this measurement. There is a notable lack of research on suitable neuropsychological tests for assessing executive functions across diverse autonomic regulatory states. This paper reviews the most frequently used neuropsychological instruments in this context, aiming to guide the research community towards optimal tasks and administration protocols for concurrent autonomic nervous system measurement.

The diversity of current executive function tests presents both opportunities and challenges. Some tests are better suited for simultaneous neurophysiological measurements due to their design, duration, and cognitive load, while others may interfere with or be influenced by such monitoring. This variability can lead to inconsistencies in findings and complicate result interpretation and comparison across studies of brain disorders. A significant drawback of using different tasks is the difficulty in comparing outcomes and conducting meta-analyses.

Standardizing a smaller selection of tasks with consistent protocols would enhance research reliability, facilitate robust comparisons, and improve the overall quality of meta-analyses. To achieve this standardization, it is essential to first survey and describe the current landscape of executive function assessments in conjunction with autonomic nervous system measurements.

**Objective:** This paper aims to comprehensively analyze current instruments utilized in the assessment of executive functions, elucidating their advantages, limitations, and implications for future standardization initiatives. By systematically examining the most prevalent tools for evaluating executive functions in conjunction with autonomic nervous system measurements within clinical and experimental research settings, this review seeks to provide valuable insights for enhancing methodological consistency and advancing research in this area.

**Methods:** We searched for articles published using the PubMed database with the following terms: **(neuropsychological test OR neuropsychological evaluation OR neuropsychological measure*) AND (executive functions OR EF OR executive function) AND (autonomic OR ANS OR parasympathetic OR PNS OR vagal OR “heart rate variability” OR HRV OR sympathetic OR HPA OR electrodermal OR RSA)**

Only the healthy population was chosen. There was no language restriction.

**Results:** 62 articles fulfilled all the inclusion criteria. The 5 neuropsychological tests most frequently used to evaluate executive functions **in relation to autonomic regulation** were:

1. Trail Making Test (TMT) B
2. The n-back Task including 2-back Task
3. Wisconsin Card Sorting Test
4. Stroop Test and its variants
5. Wechsler Adult Intelligence Scale (WAIS)-Working Memory Composite

The domains of executive functions most frequently assessed are: cognitive flexibility, working memory, and inhibitory control/selective attention.

**Conclusion:** These findings offer valuable insights for future research directions and the development of standardized assessment protocols for executive functions, tailored to diverse socio-demographic profiles.

## 1. Introduction

This paper addresses a significant gap in the research on the relationship between executive functions and autonomic nervous system regulation, which is crucial for advancing both clinical and experimental research. The autonomic nervous system, responsible for regulating vital physiological processes, is complex and time-consuming to measure, requiring meticulous methodologies that often limit the feasibility of concurrent cognitive and executive function assessments. Additionally, autonomic nervous system measurement becomes difficult due to subject movement, changing environment and time-consuming protocols. There is a notable scarcity of investigations into which neuropsychological tests are suitable for assessing executive functions across diverse autonomic regulatory states. This paper seeks to bridge this gap by systematically reviewing the neuropsychological instruments most frequently employed to assess executive functions in the context of different autonomic control systems. By doing so, we aim to guide the research community towards a consensus on the optimal executive functions tasks and their administration protocols for concurrent autonomic nervous system measurement. Establishing such a consensus is essential for standardizing experimental methodologies, enhancing the reliability and comparability of research findings, and deepening our understanding of the interaction of cognitive processes and autonomic regulation. This comprehensive article will ultimately facilitate more precise and integrative investigations in both clinical and experimental settings.

### 1.1 Executive functions, its classification, and its significance

Executive functions are complex cognitive abilities that enable the identification of goals, mental planning, behavior organization, and selective attention to achieve these goals. Executive functions (EFs), encompass a set of top-down mental processes essential for situations requiring focused attention. These functions come into play when relying on automatic responses or instincts would be ill-advised, insufficient, or impossible (Diamond, 2013; Miller & Cohen, 2001). Engaging in executive functions requires effort; it’s simpler to persist in current actions than to initiate change, easier to succumb to temptation than to resist it, and more straightforward to operate on “automatic pilot” than to contemplate the next course of action.

The classification of executive functions lacks consensus, and ambiguity in terminologies has been observed in the articles we have examined. The term “executive function” itself has been interchangeably used with “executive control” or “cognitive control” in some articles (Miyake et al., 2000). Nevertheless, in recent years, there has been a general consensus that three core EFs exist (Banich, 2009; Diamond, 2013): inhibition [inhibitory control, including self-control (behavioral inhibition, resisting impulses) and interference control (selective attention and cognitive inhibition)], working memory (WM), and cognitive flexibility (shifting, mental flexibility, or mental set shifting and closely linked to creativity). Combination of two or more of these EFs provides ground for goal directed behavior. From these, higher-order EFs are built such as reasoning, problem solving, planning and decision making.

#### 1.1.1 Working memory

Working memory is defined as a system of temporary storage and manipulation of information or working with information no longer perceptually present (Baddeley & Hitch, 1994; Chai et al., 2018). There are two categories of working memory (WM), each differentiated by content— verbal WM and nonverbal (visual-spatial) WM. WM is crucial for performing arithmetic operations, or contemplating ideas, information, or doctrine. WM plays a key role in comprehending events that unfold chronologically, as it involves retaining information about what occurred earlier and connecting it to subsequent events. Consequently, it is indispensable for understanding written or spoken language, be it in the form of a sentence, a paragraph, or more extensive discourse. WM is vital for ordering items, making plans, reasoning, fostering creativity by connecting seemingly unrelated elements and facilitating decision-making with conceptual knowledge, including consideration of past experiences and future plans (Chai et al., 2018; Diamond, 2013).

##### 1.1.1.2 Ambiguities in nomenclature and opinions on definitions

There is a difference in short-term memory and WM, short-term memory is holding information whereas WM is holding information in mind and manipulating it (Diamond, 2013). Also, there are theories which endorse that WM supports inhibitory control and they generally need one another and co-occur. Theories of working memory encompass inhibitory control within the working memory framework (Conway & Engle, 1994). While EF researchers often view working memory as a subcomponent of EFs, some working-memory researchers use the term more broadly, putting it on the same level as EFs. Engle and Kane, for instance, define working memory as the ability to maintain selected information while inhibiting distractors and interference (Conway & Engle, 1994; Shipstead & Engle, 2013). In Baddeley’s working-memory model, the central executive functions include inhibitory control and cognitive flexibility, involving multitasking, task shifting, and selective attention and inhibition. Moreover, WM and selective, focused attention appear to be similar in many ways, including their neural basis. Simulations have shown that enhancements in working memory during development can contribute to improvements in selective attention (Awh et al., 2006; Ku, 2018). Personally, we would prefer to use the term working memory (WM) exclusively for the purpose of retaining and actively manipulating information. However, in this review article, we will use the nomenclature used by the selected manuscript authors.

#### 1.1.2 Inhibitory and interference control

Inhibitory control, a fundamental executive function, involves managing attention, behavior, thoughts, and emotions to consciously choose appropriate actions over impulses or environmental influences. This ability empowers us to break free from habitual responses and make intentional choices, offering the potential for change and preventing regrettable decisions.

The two subcomponents of inhibitory control are self-control or response inhibition, and interference control. Self-control, within the realm of inhibitory control, encompasses the regulation of both behavior and emotions to guide purposeful actions (Diamond, 2012; Tiego et al., 2018). It involves resisting temptations and avoiding impulsive reactions. Additionally, self-control includes the discipline to stay focused on a task despite distractions, complete it despite temptations to quit, and resist the allure of more engaging activities. The subcomponent of interference control pertains to the regulation of attention and cognition. At the level of attention, it involves and referred as selective attention, resisting distractions in the environment, and maintaining focus, as required in noisy settings (Grinspun et al., 2020). In terms of cognitive inhibition, it involves averting internal distractions, such as extraneous or unwanted thoughts, and preventing mind-wandering and drifting (Kam & Handy, 2014; Keulers & Jonkman, 2019). Cognitive inhibition is suppressing prepotent mental representations. Cognitive inhibition leans to adhere more with WM measures than with measures of other types of inhibition.

##### 1.1.2.1 Ambiguities in nomenclature and opinions on definitions

An attention-grabbing stimulus, like visual motion or a loud noise, captures our attention involuntarily. That is called exogenous, bottom-up, automatic, stimulus-driven, or involuntary attention and is driven by properties of stimuli themselves (Nguyen et al., 2020; Theeuwes, 2010). Inhibitory control of attention allows us to selectively concentrate on our chosen focus while suppressing attention to other stimuli. Besides being called selective or focused attention, this has been termed attentional control or attentional inhibition, endogenous, top-down, active, goal-driven, voluntary, volitional, or executive attention (Matusz et al., 2018; Nguyen et al., 2020; Theeuwes, 2010).

#### 1.1.3 Cognitive flexibility

The third core executive function is cognitive flexibility, is also commonly known as set-shifting or mental flexibility. One component of cognitive flexibility involves seamlessly transitioning between different tasks or mindsets and includes the ability to change spatial or interpersonal perspectives, requiring the inhibition of previous viewpoints and activation of new ones. Thus, very often this process involves both inhibitory control and working memory (Dajani & Uddin, 2015). Another facet of cognitive flexibility is altering how we approach a problem, encouraging thinking outside the box. For instance, if a particular solution proves ineffective, can we devise a novel approach that wasn’t previously considered? Cognitive flexibility shares common ground with creativity, task switching, and set shifting, representing the opposite of cognitive rigidity (Dajani & Uddin, 2015; Uddin, 2021).

#### 1.1.4 Derivation of higher-order executive functions

From these three core EFs, higher-order EFs are built such as reasoning, problem solving, decision making, planning and overall goal directed behavior (Colautti et al., 2022; García-Molina et al., 2010). Few authors consider fluid intelligence as a higher-order executive function (Dang et al., 2014; García-Molina et al., 2010; Van Aken et al., 2016). Reasoning and problem-solving constitute the essence of fluid intelligence, with an exceptionally high correlation between working memory and fluid intelligence (Dang et al., 2014; García-Molina et al., 2010). Relational reasoning or logical reasoning are also built upon these three EFs (Dang et al., 2014; García-Molina et al., 2010; Van Aken et al., 2016).

#### 1.1.5 Significance and evaluation of executive functions

Executive functions are important tools that provides humans control over their environment. EFs are crucial skills for both mental and physical well-being and are foundational for overall well-being and success. Beyond their impact on mental and physical health, they are pivotal in professional contexts, enhancing problem-solving and adaptability. In relationships, EFs contribute to effective communication and empathy. They play a crucial role in emotional regulation and resilience, aiding in stress management. EFs are essential for goal achievement, fostering independence and autonomy. Additionally, they support adaptability, influence academic success, and contribute to healthier lifestyle choices. In essence, EFs play a multifaceted role in shaping successful individuals (Colautti et al., 2022; Gallant, 2016).

Executive dysfunction, a prevalent symptom, manifests across a diverse spectrum of psychological and psychiatric disorders, as well as in individuals with neurological damage. This impairment in executive functions, encompassing cognitive processes like decision-making, problem-solving, and impulse control, serves as a common denominator in various clinical conditions. From psychiatric disorders such as ADHD and depression to neurological conditions like traumatic brain injury or dementia, executive dysfunction often emerges as a prominent feature (Martínez et al., 2016; Song et al., 2020).

#### 1.1.6 Measurement of executive functions and challenges in selecting neuropsychological test

Measurement and evaluation of these executive functions provides an insight into neurophysiology of behavioral, cognitive, emotional, and psychological realm. Quantifying executive functions allows for the identification of patterns in behavior, cognitive processes, emotional regulation, and psychological well-being, enabling targeted interventions, tailored educational strategies, and personalized approaches to support individuals in optimizing their executive function skills. Thus, neuropsychological tests and techniques to measure these executive functions are equally important.

The abundance of neuropsychological tests for measuring executive functions complicates the selection process. Choosing the right test becomes challenging due to the diverse range of assessments available, each targeting specific aspects of executive functions. Researchers and practitioners must carefully consider factors such as focus, sensitivity, and reliability to make informed decisions about the most appropriate instrument for their specific needs. This highlights the importance of a thoughtful approach in selecting assessment tools to ensure accurate evaluations of executive functions.

The task of selecting the right neuropsychological test for measuring executive functions becomes even more challenging when considering the autonomic regulation factor within the nervous system. The integration of autonomic regulation introduces an additional layer of complexity to the EF task selection process. The interaction between executive functions and autonomic regulation necessitates a nuanced understanding of how these factors interact, further emphasizing the importance of careful consideration and strategic decision-making when choosing assessment tools. This dual consideration underscores the intricacies involved in accurately evaluating executive functions within the broader context of autonomic functioning. Through this review we are trying to make this selection process easier.

### 1.2 Autonomic nervous system and its control mechanisms

The autonomic nervous system (ANS) is a component of the peripheral nervous system that controls and coordinates involuntary physiologic processes including heart rate, blood pressure, respiration, body metabolism, digestion, body temperature, sexual arousal, and the control of organ function. The ANS operates automatically and unconsciously, allowing the body to adapt to different situations without conscious effort. It contains three anatomically distinct divisions: sympathetic, parasympathetic, and enteric. The ANS responds after receiving information from the body and external environment, by stimulating body processes, usually through the sympathetic division, or inhibiting them, usually through the parasympathetic division. Many organs are controlled primarily by either the sympathetic or the parasympathetic division. Sometimes the two divisions have opposite effects on the same organ. Overall, the two divisions work together to ensure that the body responds appropriately to different situations. The autonomic nervous system plays a crucial role in maintaining homeostasis, ensuring that the body’s internal environment remains stable despite external changes (Gaudet et al., 2013; Waxenbaum et al., 2021).

The control mechanisms of the ANS are rooted in the central nervous system (CNS), involving various brain regions including the prefrontal cortex. Although the ANS primarily operates independently to manage involuntary physiological processes, it is guided and modulated by specific regions in the CNS to ensure appropriate responses to internal and external stimuli (Thayer & Lane, 2000). Key structures involved include the hypothalamus, brainstem (particularly the medulla oblongata), and the spinal cord (Taylor et al., 2010). The hypothalamus plays a crucial role in integrating autonomic functions by receiving inputs from higher brain regions and peripheral sensors, and subsequently coordinating responses through the sympathetic and parasympathetic divisions to maintain homeostasis. It sends signals to the ANS to adjust functions such as heart rate, blood pressure, body temperature, and fluid balance. The brainstem contains essential autonomic centers that directly manage cardiovascular, respiratory, and digestive functions. The spinal cord provides autonomic pathways that connect the CNS to the peripheral nervous system. Preganglionic neurons originate in the spinal cord and travel to autonomic ganglia, where they synapse with postganglionic neurons that innervate target organs. The limbic system, which includes structures such as the amygdala and hippocampus, influences the ANS by mediating emotional responses. For example, stress or fear can activate the sympathetic nervous system, leading to the “fight or flight” response. While the cerebral cortex, particularly the prefrontal cortex, does not directly control autonomic functions, it can influence them through its connections with the hypothalamus and limbic system. Thus, conscious thoughts and emotional states can affect heart rate and digestion. This centralized control ensures a coherent and adaptive response to internal and external stimuli, maintaining overall physiological balance (Thayer et al., 2009; Thayer & Lane, 2000).

The sympathetic nervous system (SNS) and the parasympathetic nervous system (PNS) both comprise afferent and efferent fibers, offering sensory input and motor output, respectively, to the CNS. Typically, the motor pathways of SNS and PNS involve a two-neuron series: a preganglionic neuron with a cell body in the CNS and a postganglionic neuron with a cell body in the periphery, which then innervates target tissues. The enteric nervous system (ENS) is an extensive, web-like structure that can function independently of the rest of the nervous system (Waxenbaum et al., 2021). With over 100 million neurons displaying more than 15 morphologies, surpassing the combined total of all other peripheral ganglia, the ENS primarily oversees the regulation of digestive processes.

Activation of the SNS induces a state of heightened overall activity and attention, known as the “fight or flight” response. This results in increased blood pressure, elevated heart rate, initiation of glycogenolysis, and halting of gastrointestinal peristalsis, among other effects (Porges, 2009). The SNS innervates nearly every tissue in the body. On the other hand, the PNS facilitates “rest and digest” processes, leading to a decrease in heart rate and blood pressure, resumption of gastrointestinal peristalsis and digestion, and other calming effects. The PNS innervates specific areas, including the head, heart, lungs, viscera, and external genitalia. Notably, it is less extensive than the SNS, lacking significant innervation in much of the musculoskeletal system and skin (McCorry, 2007). The enteric nervous system (ENS) comprises reflex pathways that govern digestive functions, including muscle contraction, relaxation, secretion, absorption, and blood flow.

In terms of neurotransmitters, both presynaptic neurons of the SNS and PNS use acetylcholine (ACh). Postsynaptic sympathetic neurons typically release norepinephrine (NE) as their effector transmitter to act on target tissues, while postsynaptic parasympathetic neurons consistently utilize ACh. Enteric neurons employ various major neurotransmitters such as ACh, nitrous oxide, and serotonin (Tindle & Tadi, 2022).

#### 1.2.1 Central autonomic network: neuroanatomy and functional significance

The Central Autonomic Network (CAN) is a complex neural network crucial for regulating and coordinating autonomic functions in humans, situated within the CNS and encompassing key structures such as the hypothalamus, brainstem nuclei like the nucleus tractus solitarius (NTS) and dorsal motor nucleus of the vagus (DMV), limbic system components including the amygdala and insula, and the anterior cingulate cortex (ACC). Although traditionally not considered a core component of the CAN, the prefrontal cortex also contributes to autonomic modulation through connections with subcortical structures and regulatory pathways (Thayer et al., 2009; Thayer & Lane, 2000). The hypothalamus, a pivotal node in the CAN, integrates afferent inputs from higher brain centers, sensory pathways conveying visceral information, and external stimuli to modulate autonomic output, thereby orchestrating adaptive responses via the ANS. This dynamic interaction between the CAN and ANS regulates physiological parameters such as cardiovascular activity, respiratory rate, gastrointestinal motility, and thermoregulation, while also significantly contributing to emotional regulation, stress responsiveness, and homeostatic maintenance through complex connectivity and functional interactions with limbic structures.

The CAN acts as the master regulator of ANS activity, adjusting the balance between sympathetic and parasympathetic influences based on internal cues and environmental demands. For instance, in response to stress or danger, the hypothalamus triggers sympathetic activation within the CAN, leading to heightened physiological arousal such as increased heart rate and dilated pupils. Conversely, during restful periods, parasympathetic dominance slows heart rate, aids digestion, and conserves energy (Porges, 2009). Adding complexity to autonomic regulation, the prefrontal cortex is known for its role in executive functions and closely interacts with the CAN and ANS, exerting top-down control over autonomic responses based on cognitive appraisal and emotional states.

#### 1.2.2 Prefrontal cortex and executive functions

The prefrontal cortex (PFC) stands as a central hub in the control and coordination of executive functions, governing a spectrum of cognitive processes essential for reasoning, analysis, goal-directed behavior and decision-making. This region, located in the frontal region of the brain, is intricately involved in the regulation of attention, working memory, cognitive flexibility, and inhibitory control—key components of executive function (Alvarez & Emory, 2006). As the neural epicenter of higher-order cognitive abilities, the PFC integrates information from various brain regions and coordinates their harmonious functioning to facilitate adaptive responses to the environment. Damage or dysfunction in the PFC has been consistently associated with deficits in executive function, underscoring its pivotal role in the intricate neural network that underlies our capacity for complex thought and goal-oriented actions (Funahashi & Andreau, 2013; Lunt et al., 2012).

##### 1.2.2.1 Prefrontal ensemble: Brain Regions Coordinates for Executive Control

In the PFC neural ensemble, distinct regions specialize in specific executive functions, like inhibition, goal monitoring, cognitive switching and reward-driven decision-making. Different studies have demonstrated that different subregions of the PFC are associated with specific aspects of executive function, forming a sophisticated network that enables seamless coordination of these cognitive processes.

The dorsolateral PFC is implicated in working memory and cognitive flexibility, the cognitive control center, reigns in impulsive urges and guides focused thinking. It’s like the brain’s “brakes,” enabling us to resist distractions and navigate complex tasks (Henri-Bhargava et al., 2018; Jones & Graff-Radford, 2021). Damage here can lead to impulsivity and planning difficulties (S. Kim & Lee, 2011).

The medial PFC acts as the goal monitor, constantly updating our progress and adjusting strategies. It is like our internal GPS, ensuring we stay on track towards our objectives. Dysfunction in this area is linked to conditions where individuals struggle with self-monitoring and goal-directed behavior (Jones & Graff-Radford, 2021; Yuan & Raz, 2014).

The anterior cingulate cortex is crucial for attentional control, directing cognitive resources to relevant tasks. Additionally, it plays a key role in error monitoring, detecting and processing discrepancies between intended actions and outcomes (Henri-Bhargava et al., 2018; Jones & Graff-Radford, 2021). Dysfunction in this region may lead to attention deficits and difficulties in error detection (Baird et al., 2006; Jones & Graff-Radford, 2021; MacDonald et al., 2000).

The ventral PFC (vPFC) is pivotal in emotional regulation, social cognition, and reward-based decision-making. It blends logic with emotion, adding a “want” to the cognitive “should.” It plays a crucial role in weighing long-term rewards against immediate desires, guiding us towards advantageous choices. It integrates emotional information into choices, facilitating goal-aligned decisions (Jones & Graff-Radford, 2021; Miller & Cohen, 2001). Dysfunction in vPFC can lead to deficits in emotional regulation and impairments in decision-making tasks with emotional components and can lead to impulsive decisions driven by short-term gratification (Jones & Graff-Radford, 2021).

The connectivity and interactions within the PFC allow for the integration of information from diverse brain areas, providing a foundation for the cohesive execution of executive functions.

#### 1.2.3 Autonomic control and executive function

The correlation between autonomic control and executive function is a subject of significant research within the field of neuroscience. Recent studies have explored how the balance and activity of the branches of the ANS influence executive functions. The PNS response appears to be linked to optimal executive function. Research suggests that a well-regulated PNS is associated with improved cognitive performance, including enhanced attention and problem-solving skills (Barber et al., 2020; Porges, 2007). On the contrary, an imbalance in autonomic control, with heightened SNS activity, has been correlated with deficits in executive function (Porges, 2007). Healthy cognitive aging is linked to the PNS, while heightened cognitive decline is associated with excessive SNS activity (Knight et al., 2020). This connection between autonomic balance and cognitive abilities underscores the complex interaction between physiological processes and higher-order cognitive functions.

Moreover, the correlation between autonomic control and executive function has implications for understanding various psychological and psychiatric disorders. Dysregulation in autonomic balance has been observed in conditions such as anxiety, depression, and post-traumatic stress disorder (PTSD) (McGirr et al., 2010). These disorders often manifest with impairments in executive functions, highlighting a potential avenue for therapeutic interventions. Exploring the complex relationship between autonomic control and executive function not only contributes to our understanding of cognitive processes but also holds promise for developing targeted treatments for individuals facing cognitive challenges associated with autonomic dysregulation.

### 1.3 Peripheral physiological measurements in assessing Autonomic Nervous System regulation

Peripheral physiological measurements provide valuable insights into the regulation of the autonomic nervous system. Parameters such as heart rate variability, skin conductance, and blood pressure serve as non-invasive indicators of autonomic activity. Among these measurements, heart rate variability (HRV) stands out as a dynamic indicator of the balance between sympathetic and parasympathetic influences on the heart (Laborde et al., 2017; Porges, 1995). HRV reflects the variation in time intervals between successive heartbeats, offering insights into the adaptability and responsiveness of the ANS. Higher HRV is generally associated with better autonomic flexibility, while reduced variability may indicate compromised regulatory function (Goldstein et al., 2016; Lewis et al., 2012). Additionally, skin conductance, measuring the electrical conductance of the skin due to sweat gland activity, provides a real-time assessment of sympathetic arousal. This parameter is particularly useful in gauging emotional or stress responses, offering a direct link to autonomic activity.

Blood pressure is another crucial peripheral measurement contributing to the understanding of autonomic regulation. Fluctuations in blood pressure reflect the multifaceted relationship between sympathetic vasoconstriction and parasympathetic vasodilation (Jansen van Vuren et al., 2019). Monitoring blood pressure variations provides valuable information about autonomic tone and cardiovascular health. Collectively, these peripheral physiological measurements offer researchers and clinicians a comprehensive view of autonomic function, enabling the identification of abnormalities or dysregulation in the ANS that may be associated with various physiological or psychological conditions.

#### 1.3.1 Peripheral Physiological Measurements and Autonomic Nervous System Regulation in the Context of Executive Functions

Examining peripheral physiological measurements is instrumental in understanding the intricate relationship between autonomic nervous system regulation and executive functions. Parameters such as heart rate variability, skin conductance, and blood pressure can not only expend our understanding of ANS but also offer valuable insights into the dynamic interchange between the ANS and cognitive processes. Changes in these peripheral physiological measurements can indicate shifts in autonomic balance, influencing attention, emotional regulation, and decision-making—integral components of EF. For example, heightened sympathetic arousal, reflected in increased skin conductance, may be associated with challenges in inhibitory control and cognitive flexibility. HRV increases with goal-directed behavior and emotion regulation, and reduced HRV is indicative of cognitive stress (Kizakevich et al., 2019; Porges, 1995). Thayer and group are using HRV as a measurement of PNS activation (H. G. Kim et al., 2018). These measurements not only provide objective assessments of autonomic activity but also contribute to unraveling the physiological basis of cognitive and executive functions and potential implications for interventions targeting cognitive and emotional well-being.

### 1.4 Integrating Executive Functions with models of autonomic control

Polyvagal Theory, proposed by Dr. Stephen Porges, emphasizes the role of the vagus nerve in coordinating responses to environmental stimuli (Porges, 2007, 2021). The myelinated branch of the vagus, associated with social engagement, aligns with higher-order cognitive functions encompassed by EF. Optimal EF functioning may manifest in heightened cardiac vagal tone, indicative of well-regulated parasympathetic activity (Lewis et al., 2012; Porges, 2021). Conversely, challenges in EF may be associated with variations in vagal activity, influencing emotional regulation and adaptive responses. The neurovisceral integration model offers a comprehensive framework for understanding the interaction between executive functions and autonomic nervous system regulation (Thayer et al., 2009; Thayer & Lane, 2000). According to this model, the prefrontal cortex, a key player in EF, is intricately connected with the ANS, particularly through the vagus nerve. The model posits that optimal EF is associated with increased vagal tone, indicative of well-regulated parasympathetic activity. This neurovisceral integration suggests that individuals with enhanced EF may exhibit better autonomic flexibility and adaptability. Conversely, deficits in EF may coincide with dysregulation in ANS function. Exploring this dynamic relationship sheds light on the physiological basis of cognitive processes, emphasizing the importance of considering both neural and autonomic components in understanding EF. This integration emphasizes the bidirectional communication between cognitive, executive, and autonomic systems, emphasizing the potential impact of emotional and social engagement on EF as well as reverse impact of cognitive processes on bodily responses (e.g., sustained attention) (Taylor et al., 2010).

In summary, our current endeavors aim to address a critical gap in the existing research landscape which is the scarcity of investigations into which neuropsychological tests are suitable for assessing EF across diverse autonomic regulatory states. Identifying appropriate neuropsychological measures capable of capturing EF variations in experimental settings suitable for autonomic measurement bridges this research gap. Here we present a systematic review of the instruments most frequently used to assess executive functions in relation to different autonomic control systems in clinical and experimental research.

## 2. METHODS

A systematic review was conducted on December 31^st^, 2023, using the PubMed database and combining the search terms **(neuropsychological test OR neuropsychological evaluation OR neuropsychological measure*) AND (executive functions OR EF OR executive function) AND (autonomic OR ANS OR parasympathetic OR PNS OR vagal OR “heart rate variability” OR HRV OR sympathetic OR HPA OR electrodermal OR RSA)**, with no language restriction. The study included articles that fulfilled the following criteria:

1. All healthy participants.
2. Autonomic system is involved.
3. Neuropsychological tests used to assess executive functions

Cross-sectional or longitudinal studies are not excluded. The studies that fulfilled the following criteria were excluded:

1. Review studies.
2. Studies with psychiatric or neurological patients.
3. Studies with obese, HIV subjects
4. Patients with cerebral damage

To assess which executive functions tests were most used, top 5 neuropsychological tests were adopted, considering only the articles that met the inclusion and exclusion criteria.

In the landscape of executive function research, a number of terminologies and intricacies in measurement methodologies prevail. We used the following system when we faced a difficulty in categorizing the executive function and/or the neuropsychological tests:

Throughout our study, we encountered a diverse array of papers, each probing different aspects of EF and employing distinct terminology as dictated by the respective academic journals. To streamline our analytical process, we adopted a systematic approach by consolidating closely related executive functions into overarching categories. For instance, cognitive flexibility and mental flexibility, were merged into a unified category for the purposes of classification. This methodological strategy facilitated a more coherent understanding of the EF spectrum and enabled comparative analyses across studies.

A recurring challenge in our literature review pertained to the absence of precise definitions or the broad utilization of neuropsychological tests within certain studies. In cases where the parameters of a neuropsychological test were ambiguously defined or were applied in a broader context, we opted to exclude them from our tables for analysis. This deliberate omission aimed to uphold the integrity and clarity of our data presentation, ensuring that only well-defined and pertinent measures were included in our comparative analyses (Paper-1, 26).

The ambiguity surrounding the definition of executive function within some studies posed another significant consideration in our classification endeavors. In certain instances (papers-41, 44, 51), EF constructs were not explicitly delineated, prompting us to refrain from assigning specific EF domains in Tables 2 and 3. Nevertheless, in situations where the nature of the task or assessment inherently correlated with a particular EF domain, such as the Stroop task as delineated in Paper 2, we exercised discretion in allocating it to the corresponding category for classification purposes. This approach facilitated a balanced interpretation of the data, acknowledging both explicit and implicit associations between tasks and EF domains.

**Table 1.** describes the relationship between the most used neuropsychological tests and their predominant cognitive domains and summary of task description.

| **Names of EF tests** | **Predominant domain** | **Task description** |
| --- | --- | --- |
| **Stroop Test** | **Inhibitory Control** | The examination comprises three scenarios: during the initial phase, the participant is required to rapidly articulate the colors listed on a card. In the subsequent phase, they must vocalize the colors in which words like "all" and "today" are presented. Finally, in the third phase, the individual needs to identify the colors used to print words such as "yellow" and "red". |
| **Trail Making Test (TMT) form B** | **Cognitive Flexibility** | TMT Form B adds letters to test set-shifting and cognitive flexibility. In this test, participants are required to connect |
|  |  | a series of alternating numbers and letters in ascending order as quickly as possible. |
| **n-back test** | **Working Memory** | The n-back task assesses working memory by prompting participants to recall if a current stimulus matches one seen n steps earlier. It's widely used to measure cognitive abilities like memory retention and attention. |
| **Wisconsin Card Sorting Test** | **Cognitive Flexibility** | Participants sort cards based on different criteria, adapting to changing rules without explicit instructions. It's a standard tool for assessing executive function and is utilized in clinical settings to diagnose various neurological and psychiatric conditions. |
| **Wechsler Adult Intelligence Scale (WAIS)-Working Memory Composite** | **Working Memory** | The Working Memory Composite of the Wechsler Adult Intelligence Scale (WAIS) assesses an individual's ability to hold and manipulate information in mind temporarily. It includes tasks such as digit span and letter-number sequencing. This composite score provides insight into working memory capacity. |

**Table 2.**
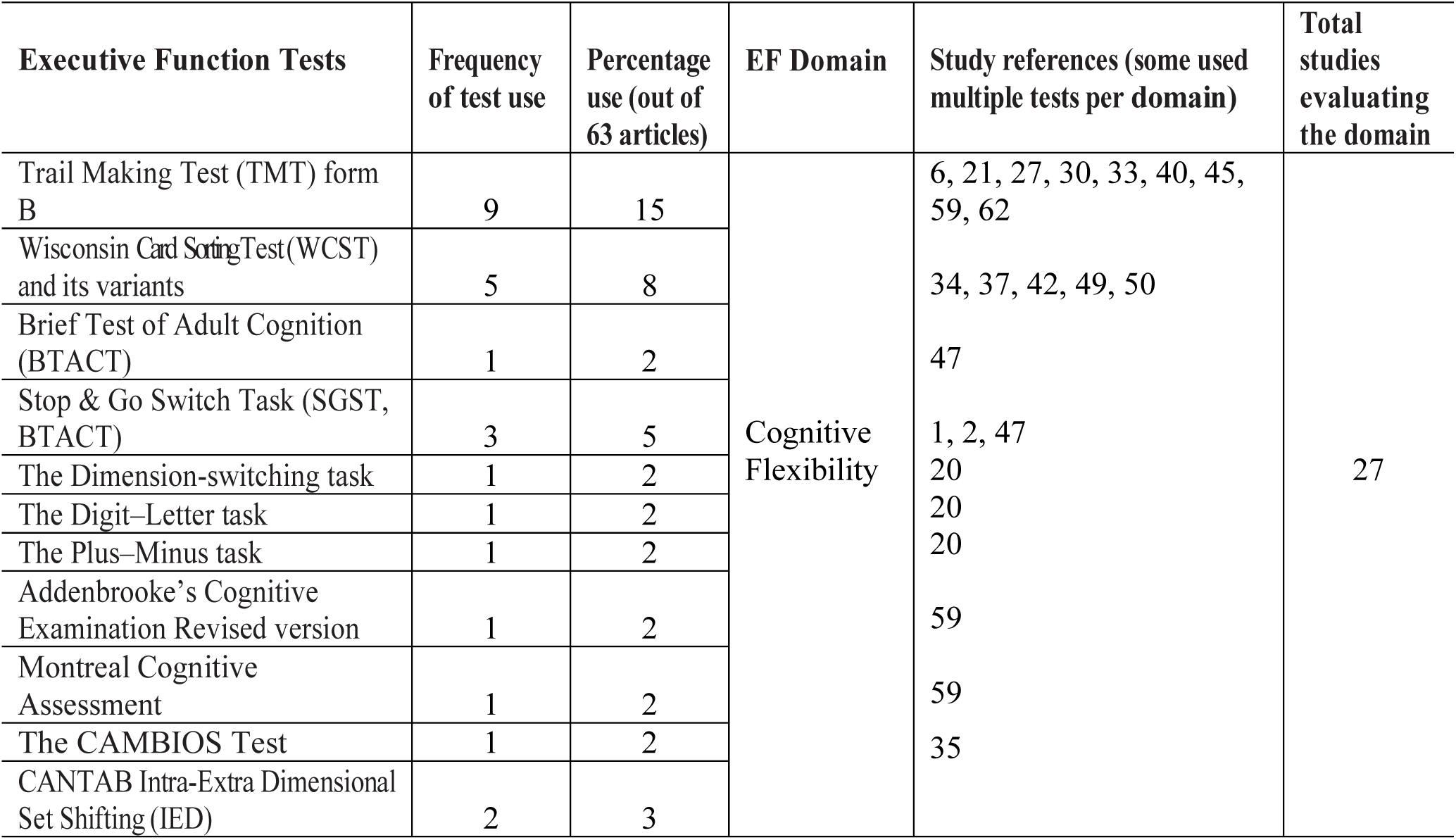
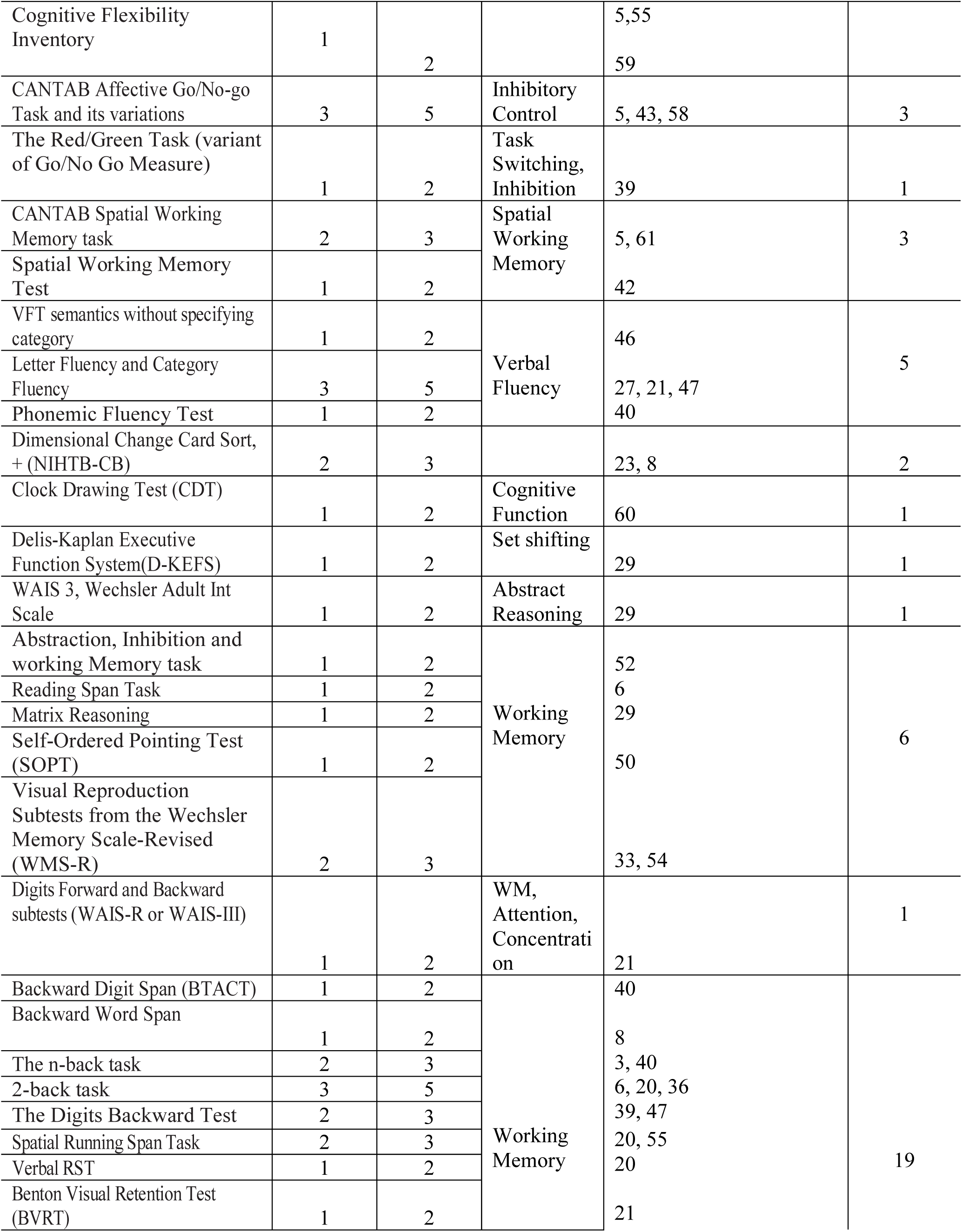
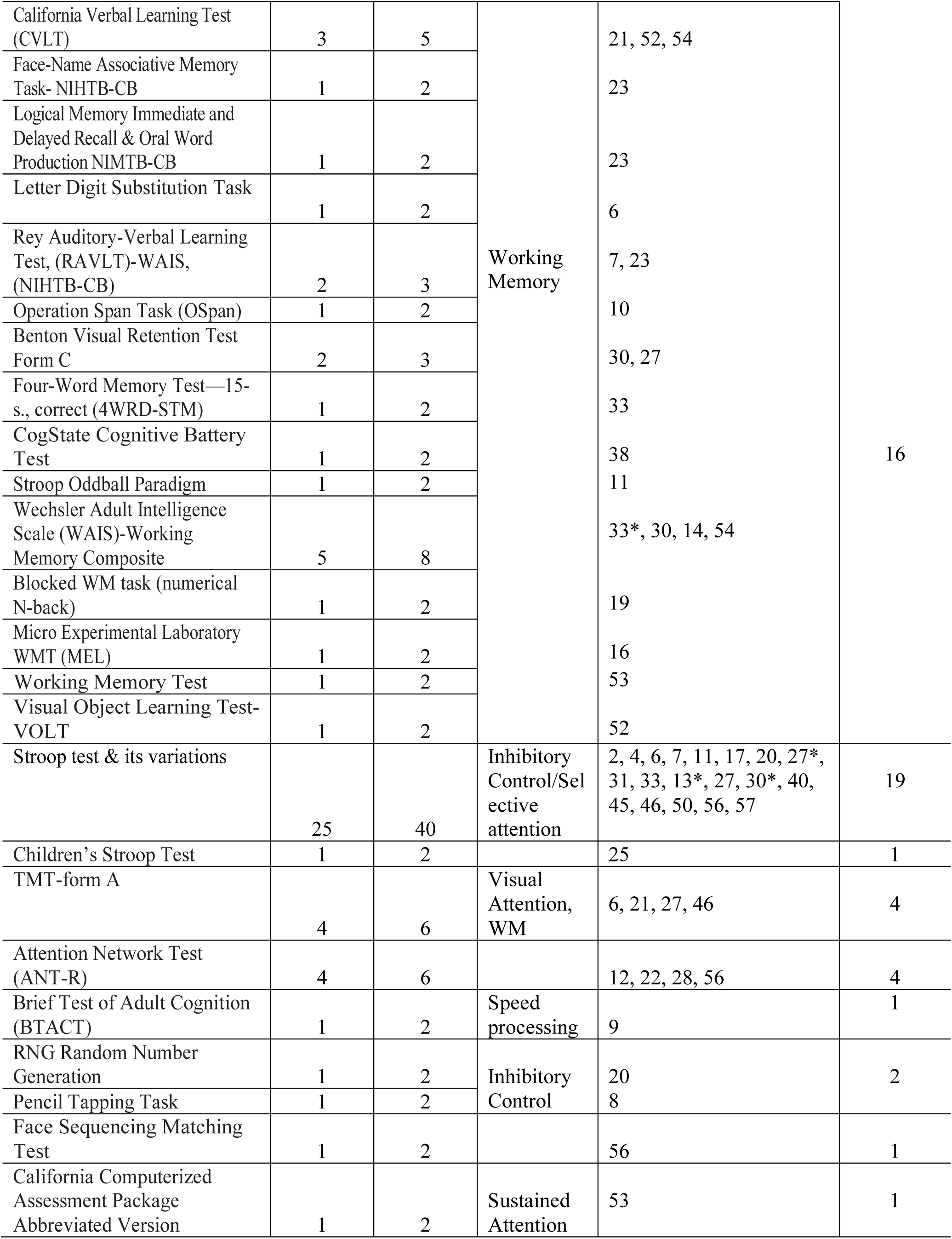
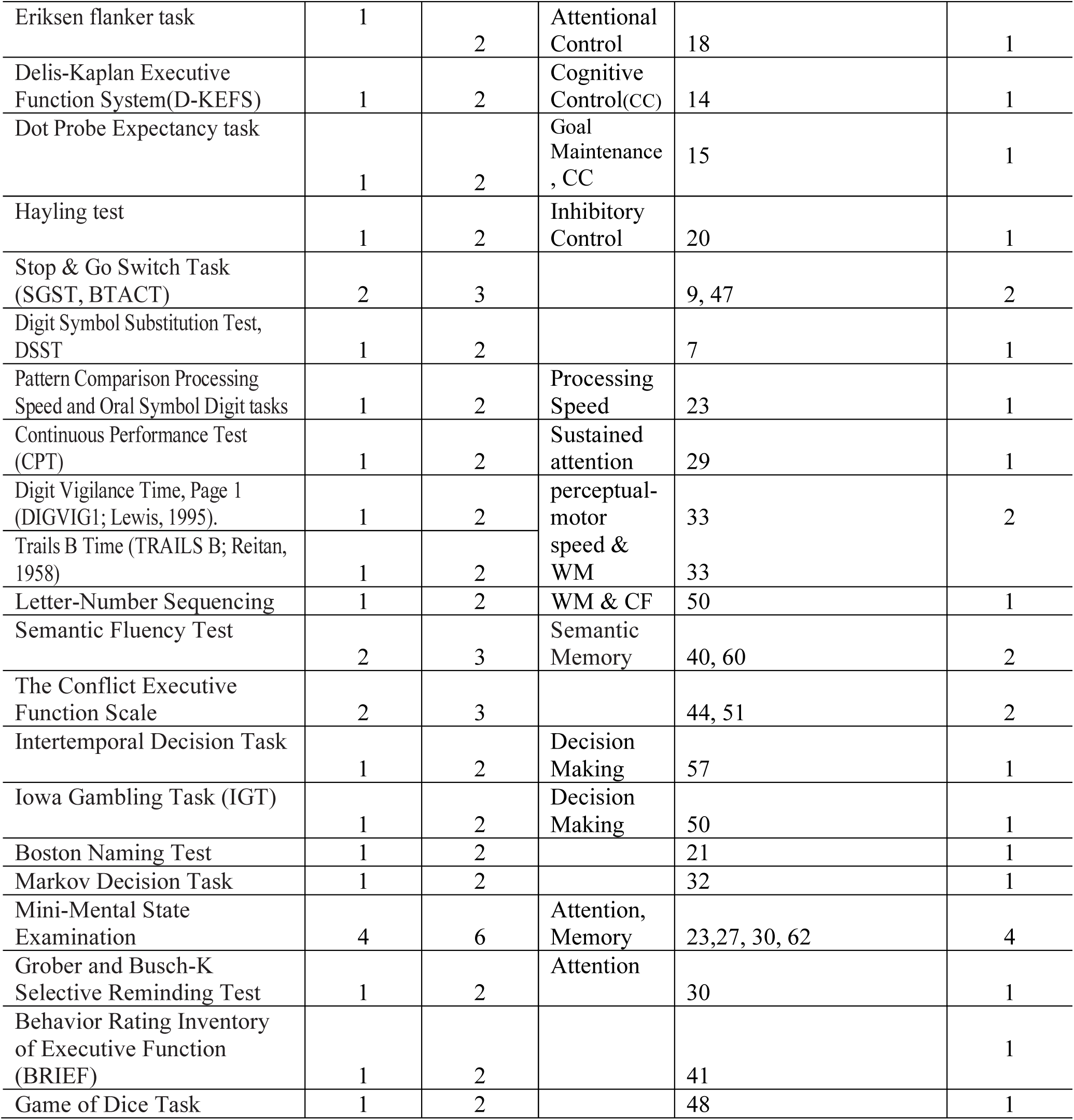
presents all the executive function tests used in the 62 articles, with their respective frequencies and percentage of usage. It also shows the number of studies by domain evaluated. In table 2, * represents either EF domain or scope of the neuropsychological test is not clearly mentioned in the paper.

## 3. RESULTS

188 articles published before 31st Dec 2023 were found in Pubmed using our search keywords. Of these, only 62 fulfilled all the inclusion criteria and 126 articles were excluded according to the exclusion criteria.

Five tests of executive function were used in equal to or more than 8% of the selected articles:

1. Trail Making Test (TMT) Form B.
2. The n-back task, including 2 back task.
3. Wisconsin Card Sorting Test.
4. Stroop test and its variants.
5. Wechsler Adult Intelligence Scale (WAIS)-Working Memory Composite

**The EF domains most frequently studied, in the decreasing order, by the researchers are, working memory, mental flexibility and inhibitory control/selective attention.**

Nineteen studies chose to use only one EF test, considering only one domain, and most studies combined two or more EF domains, as follows:

1. Only one EF domain: 19 studies
2. Combining two EF domains: 12 studies
3. Combining three EF domains: 12 studies
4. Combining four EF domains: 3 studies
5. Combining five EF domains: 4 studies
6. Combining six EF domains: 5 studies
7. Combining eight EF domains: 1 study

## 4. Discussion

The systematic review presented here identified that the tests most frequently used to assess executive functions in different autonomic regulatory systems are:

1. Trail Making Test (TMT) Form B.
2. The n-back Task including 2-back task.
3. Wisconsin Card Sorting Test.
4. Stroop Test and its variants.
5. Wechsler Adult Intelligence Scale (WAIS)-Working Memory Composite.

The EF domains most frequently studied, in the decreasing order, by the researchers are, working memory, mental flexibility and inhibitory control/selective attention.

The ambiguity in the classification of executive function domains poses a challenge in cognitive research and psychological assessments. EF encompasses a set of higher-order cognitive processes responsible for goal-directed behavior, working memory, cognitive flexibility, and self-control. However, delineating specific domains within EF, such as working memory, inhibitory control, and cognitive flexibility, can be inherently challenging due to overlapping functions and interdependencies. The lack of clear boundaries between these domains may lead to inconsistencies in measurement tools and criteria, impacting the reliability and validity of assessments. Resolving this ambiguity is crucial for refining our understanding of EF and developing more precise interventions and assessments in fields ranging from education to clinical psychology.

The nomenclature surrounding executive function introduces a layer of ambiguity in the field of cognitive and neuroscience. The terminologies used to describe specific components of EF, such as working memory, cognitive flexibility, and inhibitory control, lacks consistent standardization. Different researchers and literature may employ varied terms for similar constructs, leading to confusion and challenges in cross-study comparisons. This nomenclative ambiguity hampers the establishment of a unified framework for understanding EF, hindering effective communication and collaboration within the scientific community. Efforts to standardize the terminology associated with EF could enhance clarity and coherence in research findings, ultimately advancing our comprehension of cognitive processes and facilitating more robust discussions in the field.

As seen in Table 3, nineteen studies evaluated one EF domain and only six studies combined EF tests covering more than five domains. This can be explained by the attempt to reduce the time spent on a long neuropsychological assessment. The inclusion of numerous neuropsychological tests to assess Executive Function domains, alongside evaluations of other cognitive functions, and health parameters poses practical challenges in research settings. Conducting comprehensive assessments with an extensive battery of tests would significantly extend the duration of data collection. This prolonged testing process could lead to high dropout rates among participants, as sustained engagement might become burdensome. Moreover, the extended time requirements may deter potential participants from taking part in research studies, limiting the recruitment of a large and diverse sample. Balancing the need for thorough EF evaluations with the practical constraints of time becomes crucial, highlighting the importance of strategically selecting and optimizing neuropsychological tests to ensure meaningful data collection within reasonable timeframes for effective and feasible research outcomes.

**Table 3.** Frequency of studies by number of domains used to evaluate executive functions in various autonomic control and regulation research. **In table-3** * = one or more tests did not have clear domains while some tests did have clear tests within the paper Papers in which EF is not clear: 1, 41, 44, 51

| **Number of EF Domains Evaluated in study** | **Domains (Number of Studies)** | **Study references** |
| --- | --- | --- |
| 1 | Cognitive Control (4), Working Memory (5), Inhibition (2), Decision Making (2), Attentional Control (1), Cognitive Interference (1), Interference Control (1), Focused Attention (1), executive control (1), Cognitive Shifting (1) | 2, 3, 4, 10, 17, 18, 19, 25, 26, 32, 34, 35, 37, 43, 48, 49, 58, 59, 60* |
| 2 | Attention (7), Working Memory (5) Cognitive Control (5), Processing Speed (2), Recognition of Visual Patterns (1), Self-Control (1), Decision Making (1), Goal Maintenance (1), Interference (1), Switching (1) | 11, 12, 15, 16, 31, 36, 42, 45, 46, 53, 56, 57 |
| 3 | Working Memory (11), Mental Flexibility (5), Inhibition & Inhibitory Control (5), Verbal Learning (1), Nonverbal Learning (1), Visuospatial learning (1), Cognitive Control (5), Attention (4), Verbal Memory (1), Processing Speed (2), Short Term Memory (1), Alerting (2), Orienting (2), Memory (1), Reasoning (1) | 7*, 8, 13, 20, 22, 23*, 28, 29, 40*, 54, 55, 61 |
| 4 | Simple Reaction Time (1), Choice Reaction Time (1), Working Memory (3), Attention (1), Short-term Memory (2) & Shifting (1), Attention Shifting (1), Psychomotor Speed (1), Interference Control (1), Memory (1), Perceptual-Motor Speed (1) | 6, 33, 38 |
| 5 | Attention (3), Working Memory (5), Task Switching (1), Inhibition (2), Self-Monitoring (1), Verbal Fluency (2), Visual-spatial Learning (1), Abstraction (1), Inhibition (1), Mental Flexibility (2), Verbal Memory (1), Orientating (1), Processing Speed (1), Memory (1), Global Intellectual Efficiency (1) | 21, 30, 39, 52 |
| 6 | Working Memory (6), Verbal Fluency (2), Reasoning {&Inductive} (2), Processing Speed (4), Cognitive Control (4), Inhibition (3), Conflict Monitoring (1), Decision Making (1), Attention (2), Cognitive Flexibility (2), Emotional Processing (1), Visual Discrimination (1), Attention Switching (1), Global Intellectual Efficiency (1) | 5, 9, 27, 47, 50 |
| 8 | Set-Maintenance (1), Inhibition (1), Cognitive Control (1), Initiation and Generative Fluency (1), Verbal Fluency (1), Deign Fluency (1), Switching (1), Working Memory (1) | 14 |

In the current body of literature, there exists a notable absence of an established criterion regarding the optimal number of tests required for the evaluation of executive functions with respect to autonomic regulation. The field of cognitive science, psychology, and neuroscience particularly in the realm of executive function assessment, lacks a standardized guideline or consensus on the specific quantity of tests that should be employed for a comprehensive evaluation. This lack of clarity raises questions about the trade-off between thorough assessment and practical considerations, such as time constraints and participant burden. Researchers and clinicians are faced with the challenge of navigating this uncertainty, emphasizing the need for further investigation and dialogue within the scientific community to establish guidelines that balance the depth of assessment with the practical constraints of data collection in diverse research and clinical settings. According to the results of the present review, combining 2-3 EF domains in comprehensive batteries was the most frequent procedure adopted by 24 out of 62 articles. The choice of four or more EF domain combinations selected by only 13 articles, the is because EFs are a multidimensional construct, i.e. comprise a series of interrelated skills and high-level cognitive processing and recruit several domains in parallel, such as mental flexibility, planning, verbal fluency, inhibitory control, processing speed, and working memory. 12 studies restricted the EF assessment to two tests, usually combining a flexibility, planning or inhibitory control test with a verbal fluency test, which is quick, easy-to-apply and sensitive for discriminating people with psychiatric disorders. And 19 studies evaluated one EF domain and only six studies combined EF tests covering more than five domains.

The literature consistently demonstrates the profound influence of socioeconomic status (SES) on various aspects of human functioning, including executive functions and autonomic nervous system regulation. Studies have repeatedly shown a robust correlation between SES and executive function abilities, with individuals from higher socioeconomic backgrounds typically exhibiting superior skills in this domain compared to those from lower socioeconomic strata.

Factors such as access to quality education, early-life cognitive stimulation, and environmental resources are thought to underpin these differences, emphasizing the complex interaction between socio-environmental factors and cognitive abilities. Moreover, research has illuminated how psychosocial stressors, more prevalent in lower SES populations, can compromise cardiac parasympathetic regulation, potentially linking the stress of low SES to increased morbidity and mortality rates. The escalating burden of social stress accompanying widening SES disparities further exacerbates the impact on ANS function, highlighting the need for comprehensive understanding and interventions addressing the intricate relationship between socioeconomic circumstances and both cognitive and physiological health outcomes.

In this study, we identified the tests and specific domains commonly utilized for the assessment of executive function with respect to autonomic regulation. To the best of our knowledge, there are no other review studies in the existing literature that focus on EF testing in relation to autonomic states. The study raises concerns about the suitability of the most frequently used tests when applied to a healthy population, given the inherent variability in educational and socio-demographic levels. This variability contributes to the heterogeneity of cognitive test outcomes. Understanding the preferences of major research groups in terms of EF tests and domains can serve as a valuable guide for future research. It can aid in the development of appropriate EF assessment protocols tailored to different educational levels and socio-demographic profiles, contributing to more effective and targeted executive evaluations.

## Acknowledgment

The authors declare that I/we have no conflict of interest.

## Abbreviations

ANS: Autonomic Nervous System
EF: Executive Function
PFC: Prefrontal Cortex
HRV: Heart Rate Variability
SNS: Sympathetic Nervous System
PNS: Parasympathetic Nervous System

## Notes

### Competing Interest Statement

The authors have declared no competing interest.

